## Supplements for "Local prediction-learning in high-dimensional spaces enables neural networks to plan"

14th November 2023

#### This PDF file includes:

**1 Possible implementation of CMLs by neural networks** **2**

### 1 Possible implementation of CMLs by neural networks

The CML describes a method for online planning and problem solving on the conceptual and algorithmic level of the Marr hierarchy (Poggio, 2012). It can be implemented in various ways by biological or neuromorphic neural networks. We are describing one of them in Fig. S1. Links between mechanism which they employ and experimental data from neuroscience which support them will be discussed the next section.

This neural network implementation employs several populations of neurons: peripheral neurons that encode observations (colored blue in Fig. S1), neurons that encode actions through one-hot encoding (colored red in Fig. S1) intentions, and neurons that encode or process internal states in the high dimensional state space $S$ (colored green in Fig. S1).

The network employs in addition delay mechanisms for signals that go through inhibitory relays, which delay the signal for 1 time step. Mechanisms for implementing such delays are readily available on many neuromorphic hardware systems such as Spinnaker and the Loihi chip from Intel (Davies et al., 2021), and also on chips for in-memory computing.

One can compute utility values according to equ. 4 for all actions in parallel with the help of the learnt synaptic weights $V_{i,j}$ of the matrix $V$. A closer look shows that ideally these synaptic weights should be symmetric, like the weights in common neural network models for associative memory, such as Hopfield networks. If a reuse of the learnt weight $V_{i,j}$ from neuron j in the state representation to neuron i in the action representation as weight from i to j is not possible, one can learn the weights $W$ of the inverse connections through a simple Hebbian learning process (also referred to as perceptron learning for the case of binary postsynaptic activity) through self-supervised learning as indicated in Fig. S1b

$$\Delta W_{t+1} = \eta_w \cdot a_t (s_{t+1} - s_t)^T. \tag{S1}$$

This runs in parallel with the learning of the forward weights according to Fig. S1a. The action $a_t$ that is currently carried out provides the teaching signal (or postsynaptic activity) for the Hebbian learning process. For the case of one-hot coding of actions, that we employed for our demonstrations, this postsynaptic signal is binary. The design of the CML guarantees that a linear map $W$ exists that can carry out the desired inverse transformation, because the pseudo-inverse of the linear map $V$ is linear. Hence there exists a feasible solution for the Hebbian learning of $W$ and the Perceptron Learning Theorem of (Rosenblatt et al., 1962) guarantees then that the learning process for $W$ converges to a satisfactory solution. This separate learning of weights $W$ for the inverse connections does not reduce the resulting planning performance of CMLs in a noticeable way. In fact. we have used this method in all our demonstrations except for the ant control task, rather than reusing the learnt weights of the matrix $V$ for the inverse connections.

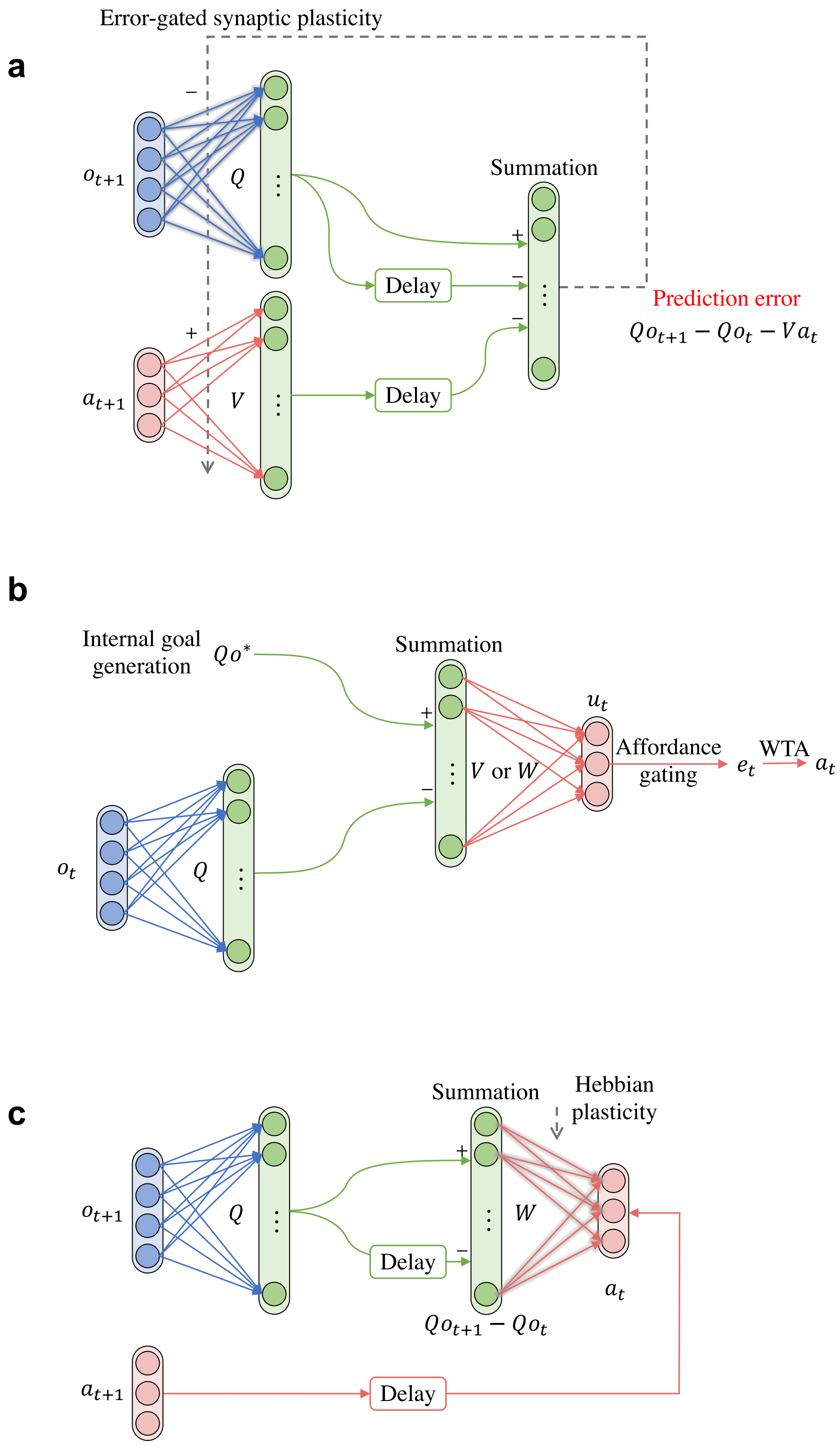
a
Error-gated synaptic plasticity
$o_{t+1}$
$Q$
$a_{t+1}$
$V$
Summation
Delay
Delay
Prediction error
$Qo_{t+1} - Qo_t - Va_t$
b
Internal goal generation
$Qo^*$
Summation
$o_t$
$Q$
$V$ or $W$
$u_t$
Affordance gating
$e_t$
WTA
$a_t$
c
Summation
Hebbian plasticity
$o_{t+1}$
$Q$
$W$
Delay
$a_t$
$Qo_{t+1} - Qo_t$
$a_{t+1}$
Delay

Figure S1: **A possible neural network implementation of the CML.** **a** Network activity during learning. The prediction error that is computed by the population of linear units on the right hand side is used to gate plasticity of the synaptic connections from populations of neurons that represent observations and actions to the populations of neurons that represent internal states of the CML. Some signals go through inhibitory interneurons (not shown here), indicated by a negative sign "-" at the corresponding synaptic connection. These signals are assumed to be delayed by 1 time step. **b** Use of the CML for planning (problem solving). A target observation $Qq*$ is given and its difference to the embedding $Qq_t$ of the current observation is computed by a population of linear neurons. Resulting utilities for all actions can be computed with the learnt weights of matrix $V$ if these weights can be assumed to be symmetric, i.e., if the learnt synaptic weight from neuron i in the action representation to neuron j in the state representation can also be used as weight for the connection from neuron j to neurons i. If this assumption is not satisfied, one can learn these weights on the side as entries of another weight matrix $W$, see the next panel. **c** If the synaptic connections whose weights are represented by the matrix $V$ can not be assumed to be symmetric, one can learn the weights from state neurons j to action neurons i (collected in a matrix $W$) on the side through self-supervised Hebbian synaptic plasticity.

### 2 Linking neuronal circuit implementations of CMLs to experimental data from neuroscience

It was shown in (Basu et al., 2021) that neural activity in the orbitofrontal cortex represents navigation goals. Such goals can be seen as a special case of the target observations $Qo^*$ that drive planning in the CML according to Principle II, see Fig. S1b. Other experimental data show that the prefrontal cortex carries out an inhibitory control over action representations, inhibiting in particular actions that would be improper in the current state. These brain mechanisms implement a form of affordance gating that is involved in action selection of the CML according to Fig. 1 c and Fig. S1b.

We propose in Fig. S1b that utilities and eligibilities are computed for all possible actions in parallel, and that the action with the largest eligibility is chosen by a WTA circuit. Such an action selection process is consistent with the experimental data of (Zagha, Ge and McCormick, 2015). The authors of this article suggest that neurons in motor cortex form a competitive circuit that regulates sensory-to-motor transformation. Furthermore, they argue that their data are “consistent with circuit models in which enhanced and suppressed neurons compete by lateral inhibition”. A model for such competition via lateral inhibition in superficial layers of cortical microcircuits, based on detailed data from the Lab of Carl Petersen, was presented in (Jonke et al., 2017).

The parallel computation of utilities for all possible actions employs the synaptic weights of the matrix W that can be learnt through self-supervised learning by Hebbian plasticity according to Fig. S1c. Alternatively, if one assumes that the synaptic connections with weights V between action representations and their embedding into the state space are symmetric, one can reuse the learnt weights of matrix V.

The learning of the matrices Q and V according to Fig. S1a employs the learning rules of eq. 2 and 3. These learning rules are special cases of the well-known Delta rule that is frequently employed in theoretical neuroscience, see Dayan and Abbott, 2005. These plasticity rules are in fact closely related to rules for synaptic plasticity that have emerged from more recent in-vivo data on synaptic plasticity, see (Magee and Grienberger, 2020; Chéreau et al., 2022) for reviews. These rules depend on presynaptic activity (modelled by the 2nd factor in eq. 2 and 3) and a gating signal (modelled by the first factor in eq. 2 and 3) that is triggered through the activity of specific populations of neurons. Like the rules of eq. 2 and 3 they do not depend on postsynaptic firing. The gating signals come according to experimental data in various forms, triggered by the firing of neurons in a variety of brain areas. For example, they come as neuromodulatory signal in the experiments of (Hong et al., 2022), as input from higher order thalamus in the experiments of (Gambino et al., 2014), as input from a hippocampal area (Doron et al., 2020), and as disinhibition of apical dendrites via the activation of VIP cells (Letzkus, Wolff and Lüthi, 2015).

The first factor in eq. 2 and 3, the gating signal, has the form of an error signal that encodes the difference between a prediction based on preceding sensory input and efferent copies of preceding motor commands and actual current sensory input as prediction target. The experimental data of (Jordan and Keller, 2020) show that there are neurons in the primary sensory cortex, so-called error neurons, that represent differences between predictions based on efferent copies of motor commands and the current sensory inputs, similar to the prediction errors that are computed on the right side of Fig. S1a. In fact, they found that different populations of pyramidal cells in layers 2/3 encode both negative and positive versions of these error signals, see Fig. 7D of (Jordan and Keller, 2020), Fig. 2A of (Keller and Mrsic-Flogel, 2018), and the graphical abstract of (Vasilevskaya et al., 2023). In fact, the recent data from (O'Toole, Oyibo and Keller, 2023) show that these error signals are computed by specific genetically encoded neuron types, thereby pointing to the involvement of genetically encoded circuitry for computing positive and negative error signals. The positive error signals are needed in the CML model to gate the plasticity of V, and the negative versions of the error signal are needed for gating the plasticity of Q, see eq. 2, and 3.

The CML model predicts that information about preceding sensory input contributes an additive offset to the responses of error neurons, provided that the current observation depends not only by the preceding motor command, but also on the preceding observation. This prediction needs to be tested in future experiments. One has already found neurons in V1 that represent a weighted sum of sensory and motor information, see (Saleem et al., 2013) and the recent review (Zhang and Xu, 2022).

Reviews such as (Wit et al., 2017) show that the psychological and behavioral role of affordances is well documented, but that there is a lack of precise knowledge how affordances are learnt and represented in neural circuits of the brain. Many brain areas, in particular PFC, appear to be involved. Also a common agreement is that affordance values arise partially from innate mechanisms for inhibitory control of action selection but that they are also subject to learning. Our model assumes that affordance values are multiplied with utility estimates for each possible action. Some forms of inhibition have been reported to have a divisive rather than subtractive impact on firing rates of pyramidal cells. Another multiplicate mechanism was recently reported in (Groschner et al., 2022). For the case of binary affordance values one can easily implement the multiplication with the affordance value by strongly inhibiting the corresponding neuron. Analog affordance values can be implemented through graded divisive inhibition. Non-binary affordance values appear in our paper only in 2 tasks: Finding shortest paths in weighted graphs and locomotion of the ant.

The neural network implementation of CML learning according to Fig. S1a and c employs also delay modules that delay signals for 1 time step in the abstract model. In biological terms this unit delay may have a duration in the range of 30 - 60 ms, since this is the time range by which signals from motor cortex to sensory cortices are delayed according to (Wang et al., 2023). According to (Mesik et al., 2019) signals from motor cortex and also numerous signals from sensory cortices reach pyramidal cells of V1 primarily through synapses on distal apical dendrites. According to (Branco, Clark and Häusser, 2010) the slow propagation of NMDA spikes in apical dendrites causes delays in the range of 50ms or more until they reach the soma. This neurophysiological mechanism is certainly able to implement delays in the range of 50ms. Other candidate neurophysiological mechanisms that could implement such delays are slow propagation of action potentials in unmyelinated axons (Debanne, 2004) and delays caused through slow activation of intermediate relay neurons. Note that all signals that are subject to a delay in Fig. S1a and c go through inhibitory interneurons before they reach their target neuron.

Altogether one sees that the learning approach of the CML is consistent with experimental data from neuroscience. In fact, it is a special case of "learning to predict", a fundamental strategy of neural networks of the brain according to (Keller and Mrsic-Flogel, 2018).

### 3 Implicit normalization of action embeddings

For online planning according to Principle II tt is desirable that all actions are mapped by the learnt weights of the matrix $V$ onto vectors in the state space that all have about the same length. Otherwise the geometry of the cognitive map would become less useful for reaching distal goals on a close-to-optimal short path as the cognitive maps shown in Fig. S2 demonstrate, and actions that are mapped onto longer vectors would also appear to have higher utility. It also ensures that the number of actions that are needed to reach a goal can be estimated with the help of the cognitive map in terms of the Euclidean distance between the start and goal state. A straightforward way to achieve that embeddings of different actions all have the same length is to normalize the columns of the matrix $V$ after each learning step:

$$V(i,j) = \frac{V(i,j)}{\sqrt{\sum_{i'} V(i',j)^2}}. \tag{S2}$$

However, it turns out that one can delete these normalization steps provided that the dimension of the state space is sufficiently high. We demonstrate this here for the task of Fig. 2 in Fig. S2a, where the planning performance and its standard deviation are shown for different dimensions of the state space. One also sees that the coefficient of variation (standard deviation divided by the mean) of the lengths of embeddings of different actions shrinks when one increases the state dimension. This is shown both before and after learning in Fig. S2b and c. Fig. S2d shows both that directions of edges in the cognitive map become more informative and that the average path length of online planning according to Principle II approaches with increasing state dimension the average shortest path length 3.302 of the best offline method, the Dijkstra algorithm, also without employing any explicit normalization.

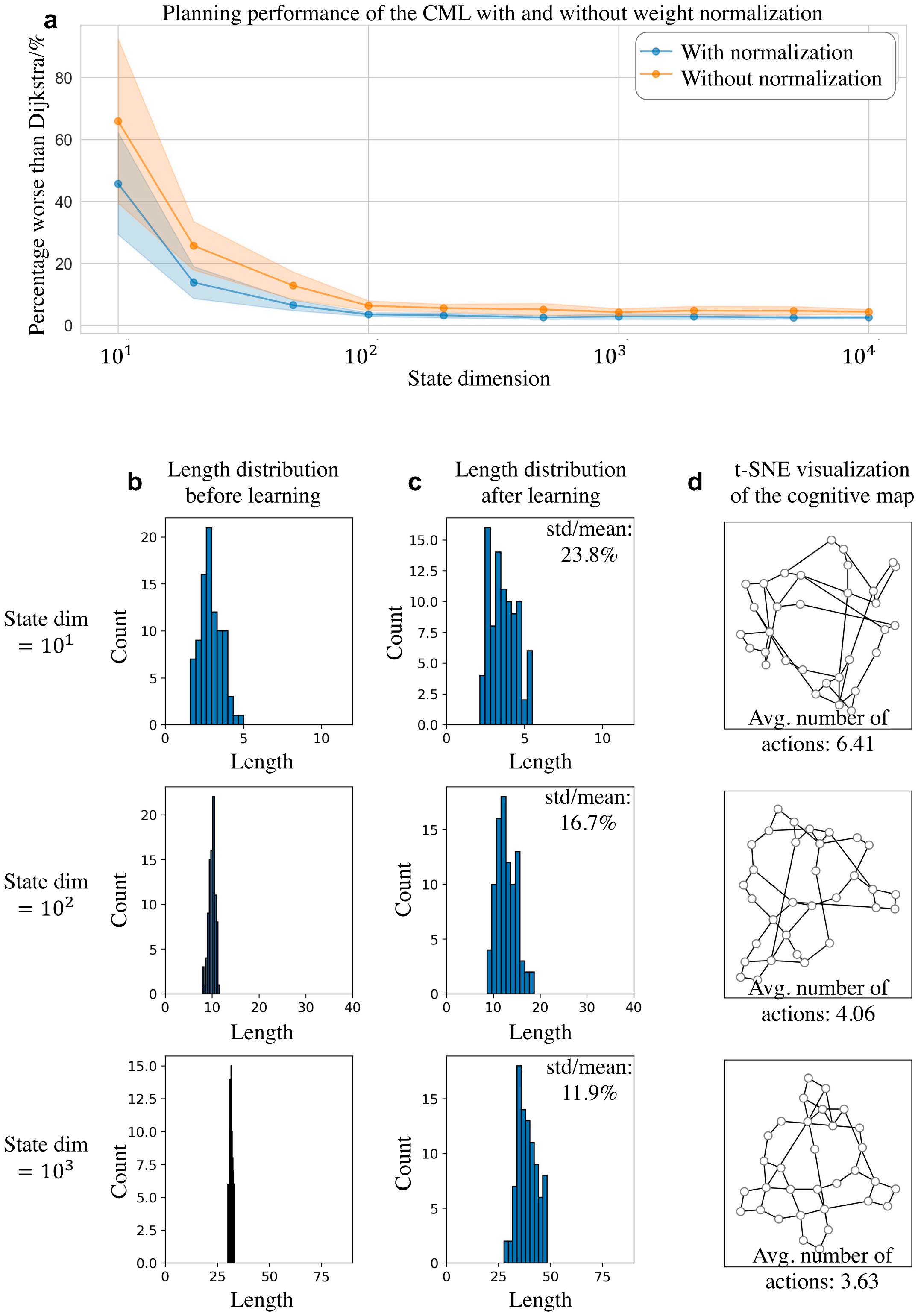
a
Planning performance of the CML with and without weight normalization
With normalization
Without normalization
Percentage worse than Dijkstra/%
State dimension
b
Length distribution before learning
c
Length distribution after learning
d
t-SNE visualization of the cognitive map
State dim = $10^1$
State dim = $10^2$
State dim = $10^3$
Count
Length
std/mean: 23.8%
std/mean: 16.7%
std/mean: 11.9%
Avg. number of actions: 6.41
Avg. number of actions: 4.06
Avg. number of actions: 3.63

Figure S2: **Impact of the dimension of the state space on the implicit normalization of action embedding vectors, and its functional impact.** **a** Comparison of the planning performance between the CML and Dijkstra (both with and without normalization) across varying state dimensions. The y-axis represents: (average CML steps/average Dijkstra steps - 1) $\times$ 100%. The solid lines (mean) and the shadowed area (standard deviation) are derived from training on 10 randomly generated graphs, testing 1000 pairs of start and goal nodes on each graph. **b** The distribution of action embeddings before learning, resulting from random initialization of $V$, for three different state dimensions. **c** After learning, there is a noticeable change in the distribution of the lengths of action embeddings. The coefficient of variation, displayed in the top-right, significantly decreases as the state dimension increases. **d** The t-SNE visualization of the learnt cognitive maps for different state dimensions shows that the Euclidean distances between nodes become more indicative of the required number of actions to reach one from the other. Additionally, the direction of each action provides with increasing state dimension better information about the set of distant goals for which this action would be useful as a first step. The average length of the path produced by the CML for different start and goal nodes is indicated at the bottom of each panel. One sees that they converge with increasing state dimension to the optimal value 3.302 that the offline Dijkstra algorithm achieves.

### 4 Further supplementary figures

We provide here supplementary figures that inform about the behavior of the CML on further tasks besides those considered in the main text. First, we show the utility values on two abstract graph environments that appear to pose particular challenges for online planning: a graph with numerous solutions, and a graph with dead ends. Then, a navigation task on a hexagonal grid, and the resulting cognitive maps are shown. One sees that the CML learns also in this spatial environment a cognitive map that captures its inherent structure, without receiving any a-priori information about the meaning of actions or the structure of the environment.

Lastly, we present for easy comparison the Dijkstra algorithm, the gold standard for offline planning. The loop structure of this algorithm is indicative of the difficulties that arise when one want to implement this planning method in neuromorphic circuits.

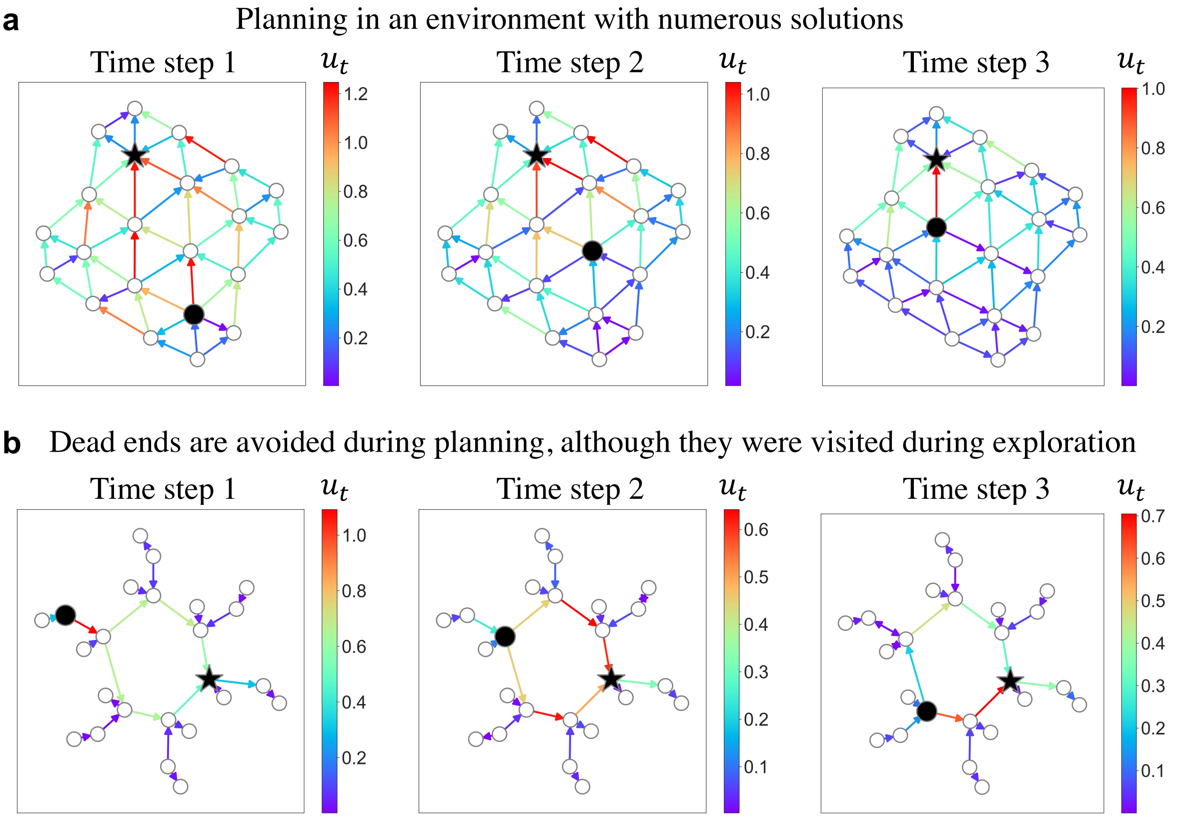

Figure S3: **Evolution of utility values $u_t$ of the CML while moving along a path, for two graphs that pose particular challenges for online planning.** Panel **a** depicts three time steps during navigation to the goal in the cognitive map for a graph that has multiple equivalent solutions to the shortest path problem. One sees that the utility values indicate at each step the best options for choosing the next action in a transparent manner. The graph was designed by hand to represent a hexagonal grid with 22 nodes, as this grid has the property allowing for multiple paths of same length from any starting node to any given target node. The shown plot was obtained by computing the t-SNE dimensionality reduction for the learnt state space of the CML. For simplicity, only positive utility values have been plotted. Note that every edge in the graph can be traversed in both directions, giving rise to two different actions in the CML. **b** The result of a corresponding experiment for a graph that was constructed to have numerous dead ends. The utility values show that the cognitive map is not confused by these multiple dead ends, although they had to be entered during learning, and always suggests next actions which avoid them. The graph was designed by hand to contain a ring consisting of 6 nodes, as well as two dead ends of different lenghts emerging from every node of the ring. For simplicity, only embeddings of actions with positive utility values have been plotted.

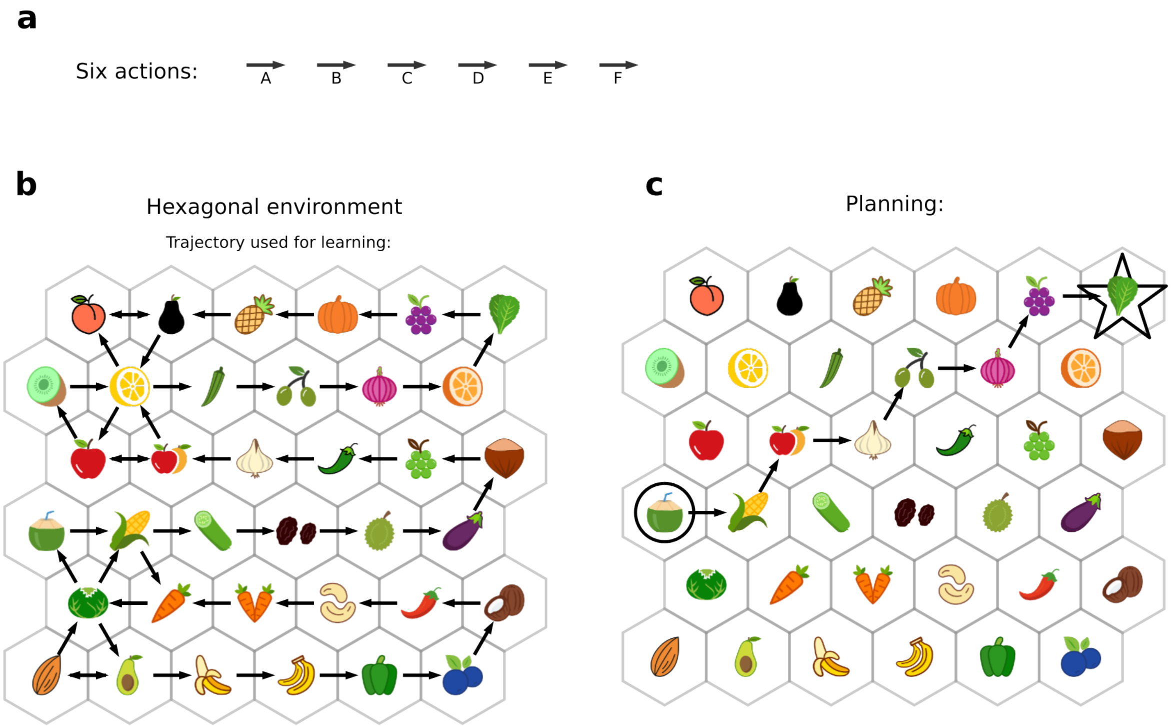

Figure S4: **Navigation task on a hexagonal grid.** The CML is challenged to produce a cognitive map for the relations between its 6 different actions and the observations it receives, analogously as in Fig. 4 for the case of a rectilinear environment. **a** Six different actions, which can be applied in any state. The CML has no prior knowledge about the meaning of them. **b** Birds-eye view of the sequence of observations which the CML received in response to a single exploration sequence with its 6 different actions. Note that most observations were experienced only once, and most of the possible combinations of observations and actions were never tried out. **c** Example for the planning capability of the CML after learning. It has to find a shortest path between the given start and goal observation. The CML produces a shortest path from start to goal, using mostly combinations of observations and actions that it never encountered during learning. Altogether this figure shows that the capability of the CML to generate cognitive maps for physical environments is not restricted to any particular geometry of the environment.

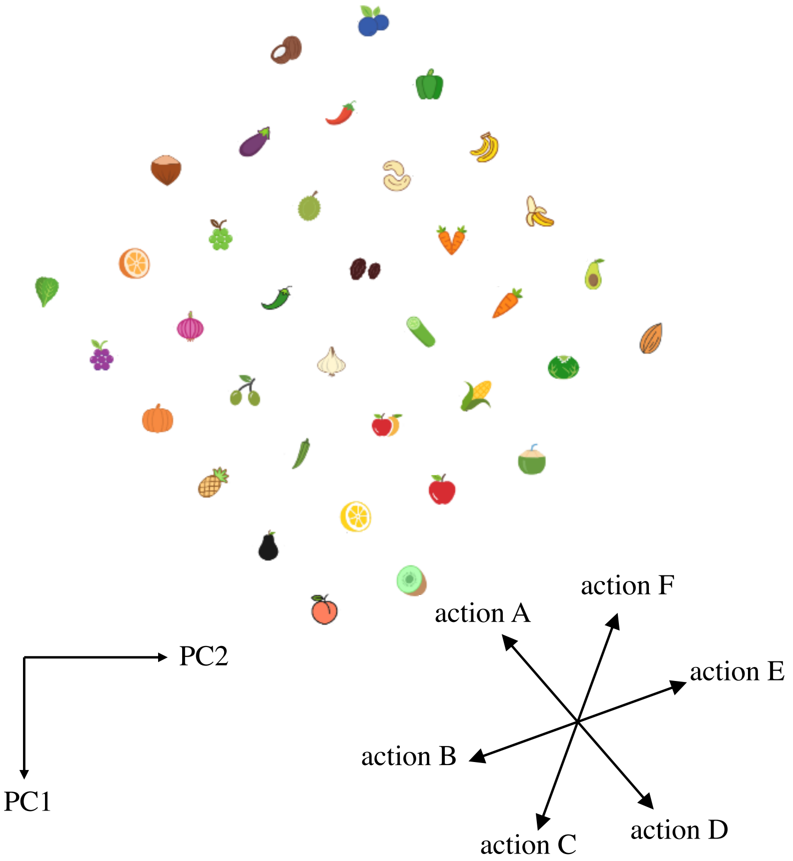

Figure S5: **The learnt cognitive map after exploration in a hexagonal environment.** Projection of the high dimensional cognitive map and the learnt action embeddings onto the first two principal components after exploration of the environment as in S4. One sees that the cognitive map has fully captured the structure of the environment from its single exploration path. This explains why the CML can produce after learning perfect solutions for any path planning challenge in this environment.

**Algorithm 1** Dijkstra algorithm for finding a shortest path in a graph

**Require:** Graph $G$
**Require:** source $S$
**Require:** target

```
for each vertex v in G.vertices do
    dist[v] ← ∞
    prev[v] ← null
    add v to Q
end for
dist[S] ← 0
while Q is not empty do
    u ← vertex in Q with min dist[u]
    remove u from Q
    for each neighbor v of u still in Q do
        alt ← dist[u] + G.edges(u, v)
        if alt < dist[v] then
            dist[v] ← alt
            prev[v] ← u
        end if
    end for
end while
S ← empty sequence                          ▷ To find the sequence to a target node
u ← target
if prev[v] is defined or u = source then
    while u is defined do
        insert u at the beginning of S
        u ← prev[u]
    end while
end if
```
